# Cortical compensation for hearing loss, but not age, in neural tracking of the fundamental frequency of the voice

**DOI:** 10.1101/2021.02.16.431374

**Authors:** Jana Van Canneyt, Jan Wouters, Tom Francart

**Author notes:** Corresponding author *Email addresses:* (Jana Van Canneyt), (Jan Wouters), (Tom Francart).

## Abstract

Auditory processing is affected by advancing age and hearing loss, but the underlying mechanisms are still unclear. We investigated the effects of age and hearing loss on temporal processing of naturalistic stimuli in the auditory system. We analysed neural phase-locking to the fundamental frequency of the voice (f0) in 54 normal-hearing and 14 hearing-impaired adults between 17 and 82 years old. We found that both subcortical and cortical neural sources contributed to the responses. Results indicated that advancing age was related to smaller responses with less cortical response contributions, consistent with an age-related decrease in neural phase-locking ability. Conversely, hearing impaired subjects displayed larger responses compared to age-matched normal hearing controls. This was due to additional cortical response contributions which were stronger for participants with more severe hearing loss. This is consistent with the recruitment of additional cortical sources for auditory processing in persons with hearing impairment.

## 1. Introduction

The auditory system, just like other systems in the human body, progressively deteriorates with advancing age. This includes loss of inner and outer hair cells, loss of spiral ganglion cells and auditory nerve fibers, as well as central processing deficits (Gordon-Salant et al., 2010). Due to these changes, many older adults report speech understanding problems, especially in noisy environments, even though they have a normal clinical audiogram. Only when the damage to the auditory system is extensive enough to prevent someone from hearing soft sounds, will hearing deficits show up in the audiogram and is the patient diagnosed with hearing loss (Wu et al., 2019). Hearing loss is one of the most common sources of disability and its prevalence is increasing (World Health Organization, 2018). Moreover, hearing loss is related to accelerated cognitive decline of older adults (Lin et al., 2013; Slade et al., 2020) and has been identified as the largest potentially preventable risk factor for dementia (Livingston et al., 2017). In this light, it is important to diagnose and treat hearing loss as early as possible. Since auditory processing is often degraded long before the audiogram indicates hearing loss, there is increasing interest for other, preferably objective measures of auditory processing.

A recent article by Anderson and Karawani (2020) reviewed various EEG-based objective measures for auditory processing in normal hearing and hearing impaired older adults. All these measures reflect temporal processing, i.e. the synchronization of the neural activity in the auditory system to the input stimulus. The reviewed measures can be divided in measures reflecting ‘subcortical’ processing (auditory brainstem responses (ABR), frequency following responses (FFR) and high frequency auditory steady-state responses (ASSR)) and measures reflecting ‘cortical’ processing (low frequency ASSR, cortical auditory evoked potentials (CAEP) and envelope tracking responses). However, it is important to note that recent studies report cortical contributions to FFRs and high-frequency ASSRs(Coffey et al., 2016, 2017; Bidelman, 2018). Thus, even though these responses are usually classified as a subcortical response, one should be careful interpreting it as a purely subcortical process.

The above-mentioned also differ in how well they approach auditory processing in daily life. As argued by Hamilton and Huth (2018) and Keidser et al. (2020), the use of natural stimuli in ecologically valid experiments is the future of auditory science. Of the cortical response measures discussed in Anderson and Karawani (2020), envelope tracking uses the most ecological valid stimulus, i.e. continuous natural speech. In contrast, CAEPs and low-frequency ASSRs require short stimuli to be repeated hundreds of times. Through encoding/decoding models (Crosse et al., 2016), envelope tracking estimates the neural processing of the speech envelope from EEG responses measured while subjects listen to a story or an audiobook. Auditory processing of these continuous meaningful speech stimuli is more relevant for daily life than traditional measures with short repetitive and unnatural stimuli (ASSRs, FFRs, CAEPs). Studies with envelope tracking have shown that older normal hearing adults have larger cortical envelope tracking responses (for speech in noise) compared to younger normal hearing adults (Presacco et al. (2016) and Decruy et al. (2019)). Therefore, cortical processing seems to be enhanced with advancing age. The effect of hearing loss on cortical processing is less clear: Decruy et al. (2020), Gillis et al. (2021) and Fuglsang et al. (2020) found enhanced cortical speech tracking responses for hearing impaired subjects compared to age-matched normal hearing subjects. In contrast, Presacco et al. (2019) found no significant effect.

From the ‘subcortical’ measures discussed in Anderson and Karawani (2020), the FFR is the most ecologically valid. It can be evoked by short speech stimuli (e.g. syllables or words), but these have to be repeated hundreds or thousands of times to increase the signal to noise ratio of the responses. For subjects of advancing age, but with normal audiogram, multiple FFR studies agree that age reduces subcortical responses to the stimulus (Anderson et al., 2012; Clinard et al., 2010; Clinard and Cotter, 2015). Once again, the effect of hearing loss is less clear: FFR studies have found that hearing loss either does not affect (Presacco et al., 2019; Roque et al., 2019), decreases (Ananthakrishnan et al., 2016; Hao et al., 2018) or enhances the response (Anderson et al., 2013a; Goossens et al., 2019).

Recently, a novel and more ecologically valid measure for ‘subcortical’ processing was developed, i.e. f0-tracking (Etard et al., 2019; Van Canneyt et al., 2020b,a; Kulasingham et al., 2020). F0-tracking is a measure for neural phase-locking to the fundamental frequency of the voice (f0), which is an important speech feature that conveys intonation, emotion and speaker characteristics. Just like the FFR, the f0 response is typically subcortically dominated with possible cortical influences. But instead of short repetitive stimuli, the f0 response is evoked by continuous speech, just like cortical envelope tracking. The f0 of the voice varies quite dramatically in natural continuous speech, and this variability is not reflected in typical FFR stimuli, like vowel and syllables. Thus, this novel measure may more accurately reflect the challenges of auditory processing in daily life than the existing measures for ‘subcortical’ processing. The analysis framework for f0-tracking is based on linear decoding/encoding models, just like envelope tracking. The technique provides information about the response strength, as well as spatio-temporal properties of the response, which may allow to disentangle cortical and subcortical response contributions.

The general goal of the present study was to investigate the effects of age and hearing loss on the auditory system using this new objective and ecologically valid measure for lower order ‘subcortical’ processing. The specific research aims of this study include: 1) Investigate the effect of age on the f0 response. From FFR studies one expects the response amplitude to decrease with age. However, a recent study by Kulasingham et al. (2020) found no significant effect of age on the f0 response. 2) Investigate the effect of hearing loss on the f0 response, with careful control for age effects. Results from FFR studies on this matter are inconclusive. 3) Disentangle the subcortical and cortical contributions to the response and how each of them is affected by age and hearing loss. This may help explain contrasting results of previous studies. 4) Study the spatial patterns of the response, i.e. how the neural activity is distributed over the scalp. Other studies have reported important changes in the distribution of the activity in the brain with age and with hearing loss (e.g. increased activity in the frontal motor cortex with hearing loss (Du et al., 2016)).

## 2. Methods

### 2.1. Dataset and subjects

The data used in this study is the same as described by Decruy et al. (2019) and Decruy et al. (2020), where the effect of age and hearing loss on cortical envelope tracking was investigated. Both of these studies were approved by the Medical Ethics Committee UZ KU Leuven/Research (S57102 and S58970). The dataset includes data from 54 normal-hearing adults (41 women, 17-82 years old) and 14 hearing impaired adults with symmetric sensorineural hearing loss who used bilateral hearing aids (8 women, 21-80 years old). Normal hearing was defined as having thresholds lower or equal to 30 dB HL for octave frequencies between 125 Hz to 4 kHz. The audiogram of the ear at which the stimulus was presented, is shown in figure 1, for each subject individually as well as the group mean. All participants were Flemish (Dutch) speaking and had no indication of cognitive impairment or learning disability.

**Figure 1:**
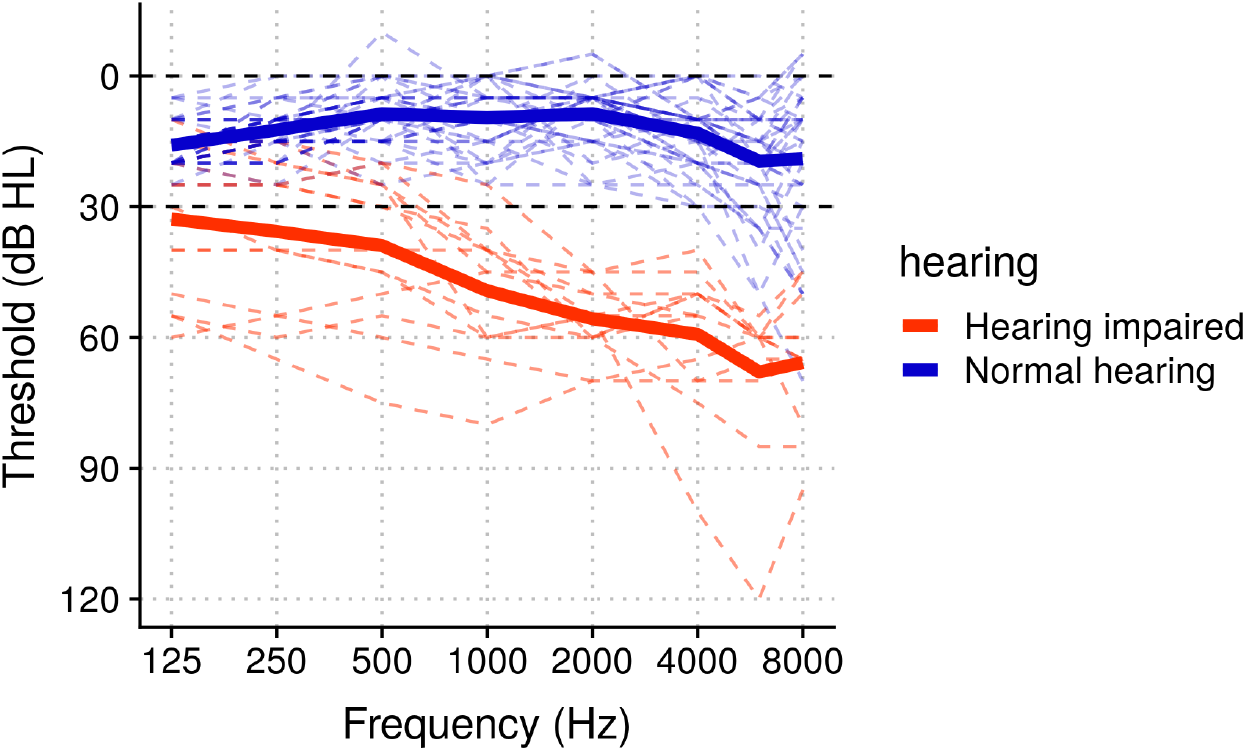
Audiogram of presentation ear. The colored dashed lines represent the pure tone thresholds of each individual. The thick lines represent the mean across individuals in the normal hearing and hearing impaired group. The black dashed lines indicate the criteria for normal hearing.

### 2.2. Stimuli

We applied the f0-tracking method to neural responses evoked by a story presented in silence. The story was 12 minutes long and written and narrated in Flemish by a male speaker (*Milan* by Stijn Vranken). The narrators voice had a median f0 of 93 Hz, and throughout the story the f0 changed with a median rate of 130 Hz/s. It was presented monaurally in the right ear (unless the left was clearly preferred on a handedness scale) through ER-3A insert phones (Etymotic Research, Inc., IL, USA). Experiment control was done using the software platform APEX (Dept. Neurosciences, KU Leuven, Francart et al. (2008)). For the hearing impaired subjects, the stimulus was amplified in a subject-specific way according to the National Acoustics Laboratory - Revised Profound algorithm (NAL-RP) (Byrne et al., 2001). This ensured that effects of hearing impairment could be studied independently of effects of audibility. The amplification was linear and implemented by filtering the stimuli with a 512-coefficient finite impulse response filter, designed based on the individual hearing thresholds. The presentation level was fixed to 55 dB A for the normal hearing participants and varied between 50 and 60 dB A for the hearing impaired participants, depending on what they reported to be most comfortable. The subjects were seated in a soundproof booth and instructed to carefully listen to the presented stimuli. The neural responses were recorded with a BioSemi ActiveTwo recording system (Amsterdam, Netherlands) with 64 active Ag/AgCl electrodes.

### 2.3. Preprocessing the EEG responses

Several preprocessing steps were performed to prepare the EEG data for f0 tracking analysis. First, the data was downsampled from a sampling frequency of 8192 Hz to 1024 Hz. Then, artefacts were removed using a multi-channel Wiener filter algorithm with delays from −3 to 3 samples included and a noise weighting factor of 1 (Somers et al., 2018). The data was re-referenced to the average of all electrodes and bandpass-filtered with a Chebyshev filter with 80 dB attenuation at 10 % outside the passband and a pass band ripple of 1 dB. The filter cut-offs, i.e. a lower cut-off at 75 Hz and a higher cut-off at 175 Hz, were based on the distribution of the f0 in the story. We also applied a notch filter to remove the artefact caused by the third harmonic of the utility frequency at 150 Hz (the other infected frequencies were not in the bandpass filter range). Finally, the unvoiced and silent sections, as determined based on the stimulus following the technique reported in Forte et al. (2017), were removed and the EEG was normalized to be zero mean with unit variance.

### 2.4. f0 tracking

The EEG responses were analysed with the recently developed f0-tracking method which is based on linear backward decoding and forward encoding models (Etard et al., 2019; Van Canneyt et al., 2020b,a). Backward modelling results in a reconstruction accuracy, which is an estimate of response strength. The results of forward modelling provide information about the spatio-temporal properties of the response. All response processing was implemented in MATLAB R2016b (The MathWorks Inc., 2016) using custom scripts and the mTRF toolbox (Crosse et al., 2016). A description of the main methods is provided here, but for details we refer to Van Canneyt et al. (2020b) and Van Canneyt et al. (2020a).

#### 2.4.1. Backward modelling

In backward linear modelling or decoding, one reconstructs a known stimulus-related feature based on a linear combination of the time-shifted data from the EEG electrodes. For f0-tracking, the feature is a waveform oscillating at the instantaneous f0 of the stimulus. As shown in our previous work, Van Canneyt et al. (2020a), an optimal f0 feature for backward modelling can be obtained by modelling the neural response to the stimulus in two steps: 1) simulating the population response in the primary auditory nerve, evoked by the stimulus, with a phenomenological model (Bruce et al., 2018) and 2) applying a low-pass filter to approximate the decreasing amplitude-frequency relation of higher level processing. The order and cut-off frequency for this low-pass filter were optimized in a data-driven way. The optimal parameters for the current dataset were equal to those for the dataset used in Van Canneyt et al. (2020a), i.e, an 8th order filter with 110 Hz cut-off frequency. This is expected as both studies used the same stimulus. The f0 feature was then filtered with the same bandpass filter that was applied to the EEG. The silent and unvoiced sections were removed from the f0 feature, after which the feature was normalized to have zero mean and a variance of 1.

The backward model was estimated by finding the linear combination of all 64 EEG channels and their time shifted versions that best approximated the f0 feature. Based on the forward modelling results, we chose to include time shifts between 0-40 ms and 0-75 ms for the normal hearing subjects and hearing-impaired subjects respectively. First, a section of the data (including minimum 2 minutes of voiced data) was set aside for testing and the model was estimated based on the remainder of the data. Regularization was done using ridge regression (Tikhonov and Arsenin, 1977; Hastie et al., 2001; Machens et al., 2004). Then, the estimated model was used to reconstruct the feature for the testing data. Finally, the reconstruction accuracy was calculated as the bootstrapped Spearman correlation between the reconstructed feature and the actual f0 feature of the test section (median over 100 index-shuffles). To validate the backward decoding results, we used a 3-fold cross-validation approach. The final reconstruction accuracy, i.e. the median correlation over the folds, is a measure for f0 respone strength. This was compared to a significance level (based on correlations with spectrally-matched noise signals) to evaluate its statistical significance (*α* = 0.05).

#### 2.4.2. Forward modelling

In forward modelling, one attempts to predict the data in each EEG channel based on a linear combination of the feature and time-lagged versions of the feature using the same ridge regression approach. In this case, time lags from −50 to 100 ms with 1/fs steps (fs = 1024 Hz) were taken into account. The weights of the forward model, also called temporal response functions (TRFs) (an average over channels as a function of time), reflect the impulse response of the auditory system, and also through topoplots, which reveal the spatial distribution of the response at a specific time lag (or the average over a range of time lags). Because the model of the auditory periphery includes compensation for frequency specific delays on the basilar membrane, using the model-based feature for forward modelling would influence the estimation of response latency. Instead, we performed the forward modelling with the ‘basic’ f0 feature used in Etard et al. (2019) and Van Canneyt et al. (2020b), which is obtained by bandpass filtering the stimulus with the same filter applied to EEG. This feature was also normalized and cut to only contain voice sections.

Because of the large degree of autocorrelation in the f0 feature, the TRFs have a periodic nature and response energy is spread in time, both in the TRFs and the topoplots. To help with interpretation, we applied a Hilbert transform when calculating the TRFs (see Etard et al. (2019)). This allows to disregard the phase and focus on amplitude variations in the TRF, but the underlying autocorrelative smearing should be kept in mind. To evaluate at which latencies the TRFs were significant, we determined a significance level (*α* = 0.05) based on forward modelling of mismatched combinations of feature and EEG data. To statistically evaluate the paired difference between two topoplots or two TRFs, a cluster-based permutation test from the mass-univariate ERP toolbox (Groppe et al., 2011) was applied. A significance level of 0.05 was used and correction for multiple comparisons is implemented within the cluster test. For more details on forward modelling and statistics, we refer to our previous work: Van Canneyt et al. (2020b).

## 3. Results

### 3.1. The effect of age

First, we investigated the effect of age on f0-tracking based on the data of the clinically normal hearing subjects only. In figure 2, the reconstruction accuracies obtained with backward modelling, estimating response strength, are presented as a function of subject age. Reconstruction accuracies ranged between 0 and 0.09 with a mean correlation across subjects of 0.035 (standard deviation = 0.025). There was a significant negative relation between age of the subject and reconstruction accuracy (r = −0.4, p = 0.003, Pearson correlation in R Core Team (2018), *α* = 0.05), indicating a reduction in f0 response strength with advancing age. In fact, many older subjects did not have significant reconstruction accuracies.

**Figure 2:**
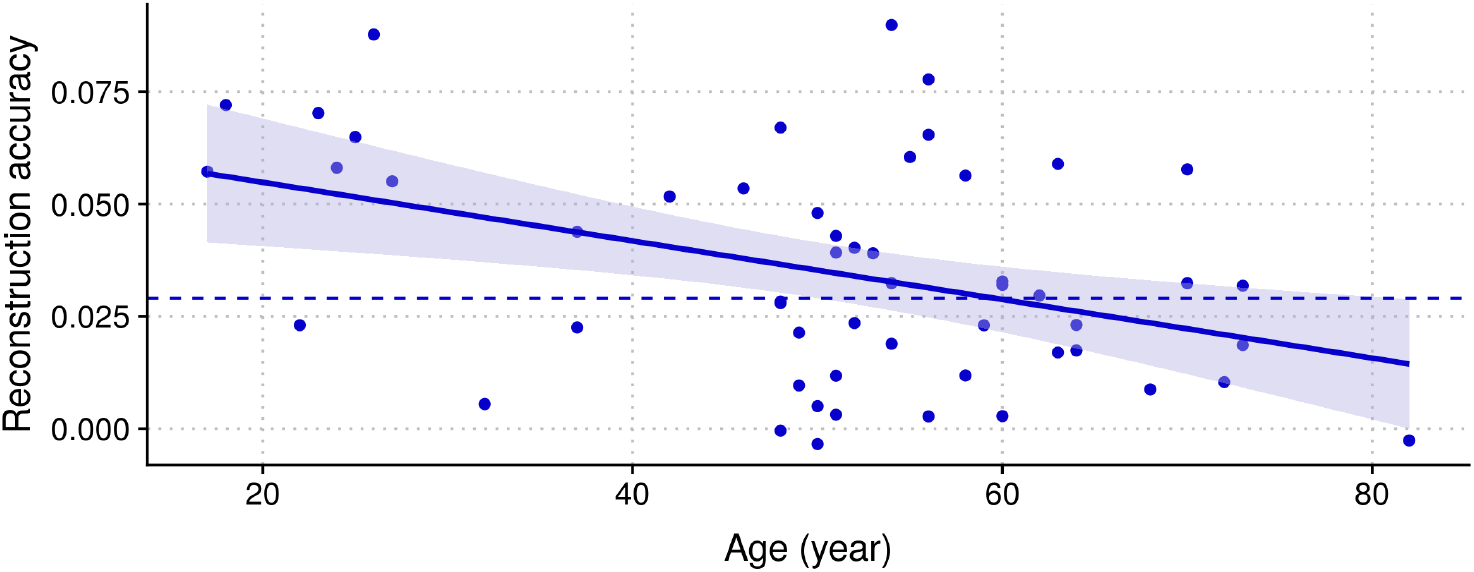
Reconstruction accuracy in function of age. The solid line presents a linear model that was fitted on the data in R. The shaded area indicates the 95 % confidence interval. The significance level for the reconstruction accuracy is indicated with a dashed line.

The spatio-temporal properties of the responses, investigated through forward modelling, are presented in figure 3. The electrode selection over which the TRFs were averaged, chosen based on the topoplots, is indicated on the figure, and includes mainly central, mastoidal and occipital electrodes. As is often the case, the TRFs vary widely in both morphology and amplitude over individuals. Therefore, we divided the subjects in three age groups and studied the average TRF in each group (see panel A). The groups were: 17-38 years old (11 subjects, mean age = 26.18, standard deviation = 6.7), 39-60 years old (31 subjects, mean age = 52.5, standard deviation = 4.5), 61-82 years old (13 subjects, mean age = 68.7, standard deviation = 5.9). The TRF was significantly different from the noise floor between 5 and 40 ms for both young and middle aged subjects (< 60 years old). For the older adults, the TRF was only significant for lags between 14.3 and 19.4 ms. Larger TRF amplitudes appear for younger subjects compared to middle-aged and older subjects in the 5 to 40 ms range. To quantify this relation, on a subject-specific level, we averaged the TRF amplitude across the 5 to 40 ms lags for each subject and correlated it with age. As presented in panel B, there was a significant negative relation between mean TRF amplitude and the age of the subject (*β* = −0.0072, df = 52, t = −3.284, p = 0.002). In panel C, the mean topoplots across six latency ranges are presented, visualising the spatial distribution of TRF activity for each of the age groups. The results indicated mostly centrally located activity which reduced in amplitude over age groups. Additionally, the topoplots of the young subject present right lateralized mastoidal activity, which is reduced in the middle-aged group and absent in the older group.

**Figure 3:**
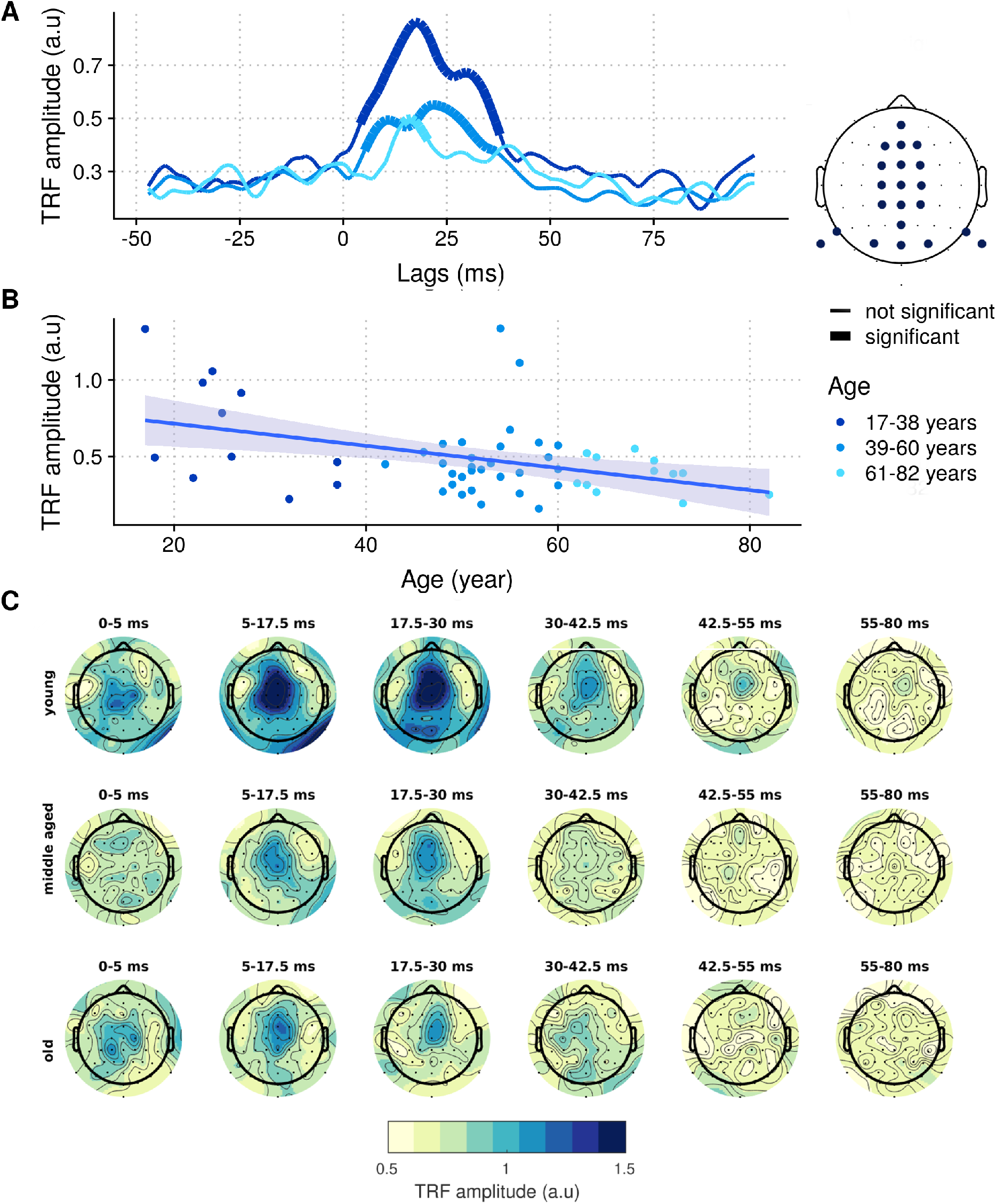
The effect of age on spatiotemporal properties of f0 tracking. **A.** Temporal response functions per age group (indicated with color). Significant sections are indicated with a thicker line. The electrode selection over which the TRFs were averaged is indicated on the head plot. **B.** Mean amplitude in the 5 to 40 ms section of the TRF per subject correlated with age. **C.** Topoplots per age group for different lags.

### 3.2. The effect of hearing loss

To study the effect of hearing loss, while controlling for the effect of age, we age-matched subjects from the normal hearing group to the 14 hearing impaired subjects (as also done by Decruy et al. (2020)). The mean age of the hearing-impaired group was 57.8 years (standard deviation = 19.9 years) and the mean age of the normal-hearing group was 57.5 years (standard deviation = 19.0 years). Panel A of figure 4 presents the reconstruction accuracies for each of these groups. As expected based on the age of the subjects, the reconstruction accuracies for the normal hearing group were small (median = 0.023) and often not significant. More surprisingly, age-matched subjects with a hearing impairment had large and mostly significant responses with a median of 0.05. A Wilcoxon rank sum test (*α* = 0.05) confirmed a significant difference in reconstruction accuracies based on hearing status (W = 144, p = 0.035). A linear model indicates that hearing impairment significantly enhanced the f0 response (*β* = −0.034, df = 25, t = −2.77, p = 0.010), even when controlling for age (*β* = −0.0007, df = 25, t = −2.32, p = 0.028).

**Figure 4:**
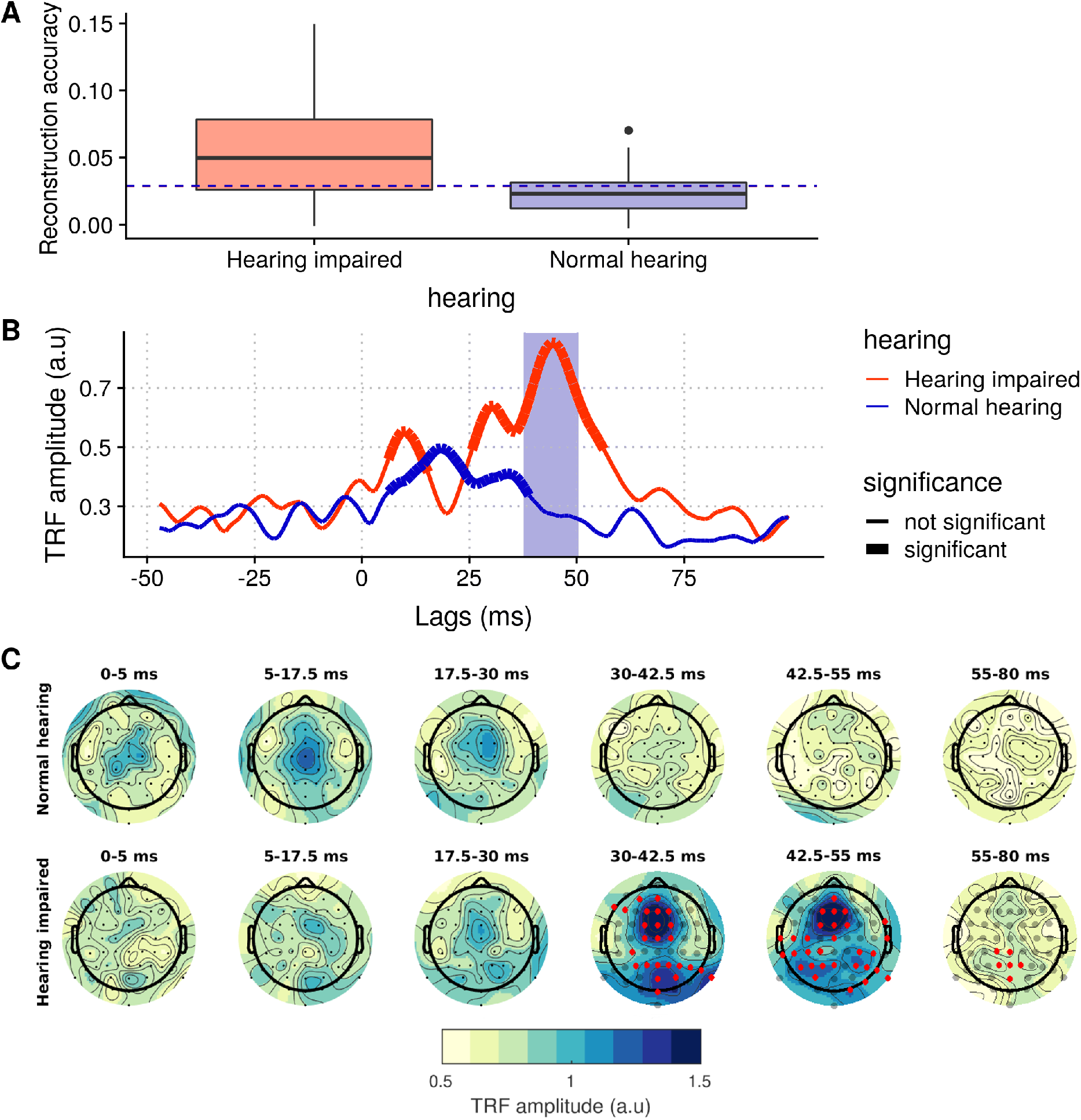
The effect of hearing loss on the f0 response. **A.** Reconstruction accuracies for age-matched normal hearing and hearing impaired subjects. The significance threshold is indicated with a dashed line. **B.** TRFs for age-matched normal hearing and hearing impaired subjects. Sections were the TRF is significantly different from noise are indicated with a thicker line. Sections were the TRFs significantly differ from each other are indicate with a purple background. **C.** Topoplots for age-matched normal hearing and hearing impaired subjects. Channels indicated with red are significantly larger in the hearing impaired subjects compared to the age-matched normal hearing subjects.

The results of forward modelling are shown in panel B and C of figure 4. The TRF analysis in panel B is based on the same electrode selection as used earlier. The TRF is significantly different from noise for latencies between 6.1 to 14.3 ms and 25.6 to 55.3 ms for the hearing impaired subjects and between 6.1 to 37.9 ms for the age-matched normal hearing subjects. Compared to the normal hearing group, the subjects with hearing loss have larger TRF amplitudes. A cluster-based permutation test from the mass-univariate ERP toolbox (Groppe et al., 2011) identified a cluster for latencies between 37.8 and 50 ms which was significantly different between the groups (p = 0.038).

In panel C, the mean topoplots across six latency ranges are visualised. In the normal hearing subjects, the majority of the response energy occurred with lags between 5 and 30 ms and this activity was mostly centrally located, as also observed in the previous section. For subjects with a hearing impairment, the majority of the response energy occurs later, between 30 and 55 ms. Those subjects present strong central activation with additional response energy distributed throughout the posterior half of the head. A cluster-based permutation test from the mass-univariate ERP toolbox (Groppe et al., 2011) was applied to statistically evaluate the paired difference between the two topoplots at each lag section. A significance level of 0.05 was used and correction for multiple comparisons (64 channels) is implemented within the cluster test. Results indicate no significant differences in the early responses (< 30 ms). However, for the later lags the responses were significantly larger in the hearing-impaired subjects compared to the age-matched normal hearing subjects across a broad channel selection. The cluster analysis identified two clusters in the 30-42.5 ms range: one frontrocentral cluster (p = 0.007: AF3, F1, F5, F7, FC1, C1, AFz, Fz, F2, FCz, FC2, Cz, C2) and one parietal cluster, which appears stronger on the right side of the head (p = 0.014: CP3, P1, P3, Pz, POz, Oz, P2, P4, P6, P8, P10, PO4). Futhermore, three significant clusters were identified in the 42.5 - 55 ms range: one central cluster (p = 0.029: F1, FC1, AFz, Fz, F2, FC2, FCz, Cz), one central-parietal cluster on the left side of the head (p = 0.010: C3, C5, T7, TP7, CP5, CP3, CP1, P1, P3, P5, PO3, Pz) and one central-parietal cluster on the right side of the head (p = 0.014: FT8, T8, CP6, CP4, P4, P6, P8, P10, PO8, O2). Finally, in the long latency range between 55 and 80 ms a small parietal cluster with significantly larger activity for hearing impaired subjects remained (p = 0.010: CP1, P1, POz, Pz, CPz, P2).

### 3.3. The effect of degree of hearing loss

To investigate whether f0 response strength was significantly related to the degree of hearing loss of the subjects, we correlated the reconstruction accuracies and mean TRF amplitude (between 30 and 55 ms) with the pure tone average (PTA) of the subjects. PTA is a measure for the degree of hearing loss, obtained by averaging pure tone audiogram thresholds for a certain frequency range, in this case 500-4000 Hz. PTAs below 25 dB HL are considered normal hearing. The results are presented in figure 5. In panel A, PTA is correlated with the reconstruction accuracies. Using linear modelling in R (version 3.6.3., R Core Team (2018), *α* = 0.05) a significant positive linear relationship was found between PTA and reconstruction accuracies (*β* = 0.0009, df = 25, t = 3.58, p = 0.001), while controlling for the age of the subjects (*β* = −0.0009, df = 25, t = −2.977, p = 0.006). In panel B, the relationship between PTA and the TRF amplitude is visualised. Again, the results indicated a significant positive relation between PTA and TRF amplitude late range (*β* = 0.009, df = 25, t = 2.98, p = 0.006), even while including the (non-significant) effect of age in the linear model (*β* = −0.006, df = 25, t = −1.81, p = 0.08).

**Figure 5:**
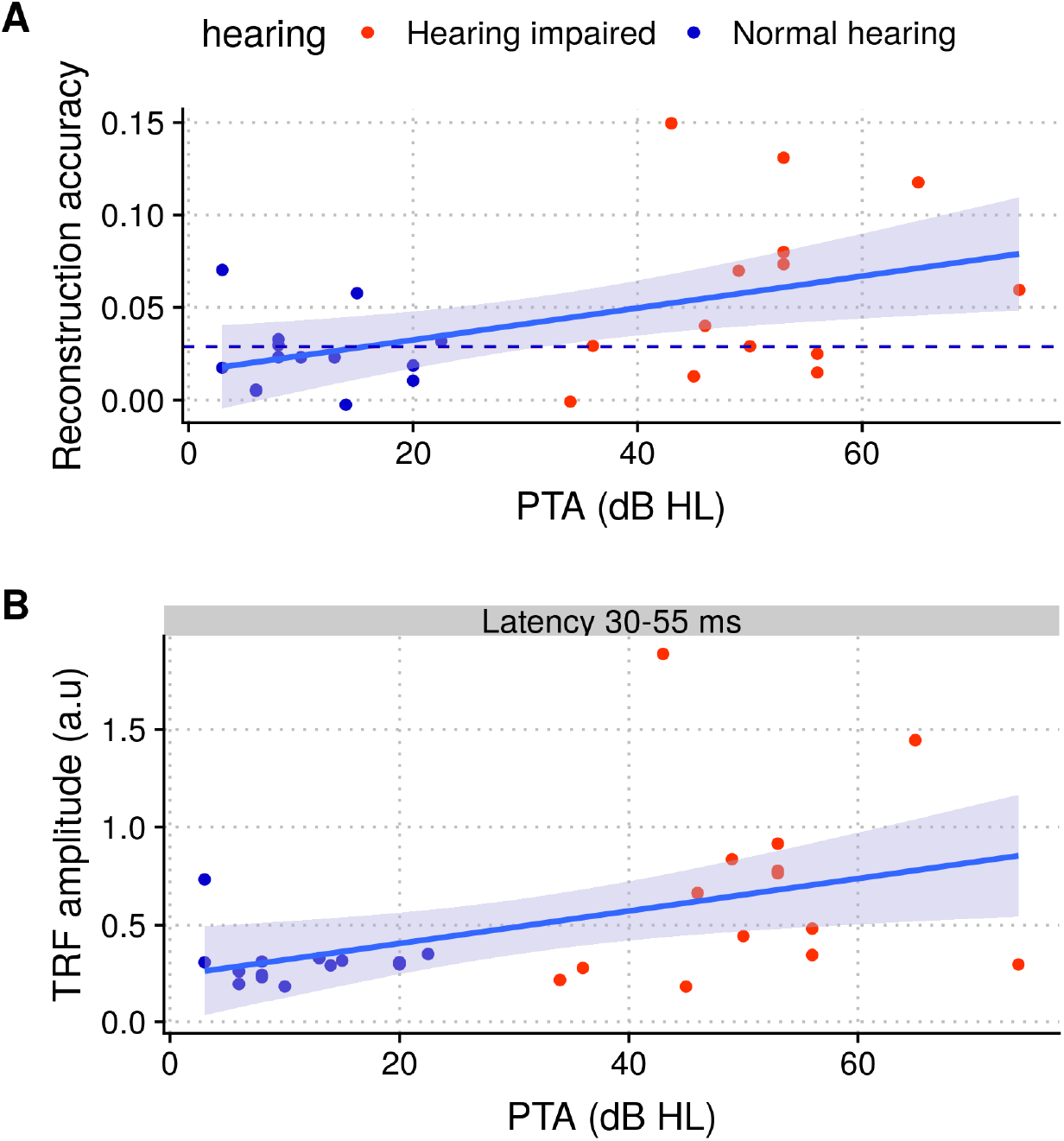
The relation between response strength and degree of hearing loss. **A.** Reconstruction accuracies correlated with PTA. The colors indicate subjects in the normal hearing group and subjects in the hearing impaired group. The line is fitted on the data using linear modelling. **B.** Mean TRF amplitude in the 30-55 ms section of the TRF per subject correlated with PTA. The latency ranges is based on the significantly different latency range between both groups indicated in figure 4.

## 4. Discussion

In this study, we investigated the effect of age and hearing loss on temporal processing in the auditory system. We employed f0 tracking, which is a novel objective measure to quantify neural phase-locking to the fundamental frequency of the voice. It uses continuous speech stimuli, which are more ecologically valid than the stimuli of other measures like the FFR and the ASSR. We analysed EEG data from both normal hearing and hearing impaired subjects in a wide age range and studied both response strength and spatio-temporal response patterns.

### 4.1. The effect of age on the f0 response

First, we investigated the effect of age on the f0 response. The results show that response strength decreased with advancing age, both in terms of reconstruction accuracies and mean TRF amplitudes. This suggests that older subjects have more difficulty with neural phase-locking at frequencies in the range of the f0. Our results are in line with the findings of multiple prior studies with FFRs, i.e. decreased responses with advancing age (Anderson et al., 2012; Clinard et al., 2010; Clinard and Cotter, 2015). In contrast, Kulasingham et al. (2020) found no significant effect of age on the f0 response. This deviant result may be explained by the fact that they used MEG to record the response, which is insensitive for radial sources, like the brainstem. As pointed out by Anderson and Karawani (2020), it is important to note that even though all subjects had clinically normal hearing, it is likely that the older adults in the group had a less pristine auditory system than the younger adults (Gordon-Salant et al., 2010). Thus, the effect of age on the f0 response is likely mediated by age-dependent factors that affect the auditory system, like anatomical changes, physiological changes and life-long noise exposure. Disentangling those factors is an interesting challenge for future research.

By calculating TRFs through forward modelling, we studied the temporal properties of the f0 response. Precise response latencies are hard to determine because of the large degree of autocorrelation in the f0 feature, which smears response energy over lags (Van Canneyt et al., 2020b). However, we can study in which latency range the response occurs. The young and middle-aged subjects displayed significant response activity with latencies between 5 and 40 ms. For older subjects the significant latencies were limited to 14 and 19 ms. The f0 response, as well as the FFR, are typically thought of as subcortical responses, because cortical neurons lack the high speed processing ability of neurons in the brainstem (Chambers et al., 2016) and cannot synchronize to the f0 when it is higher than about 150 Hz (Bidelman, 2018). However, cortical contributions can occur in normal hearing young adults when the f0 in the stimulus is low (< 150 Hz) and relatively slow-varying (Van Canneyt et al., 2020b), as is the case for the stimulus used in the present study. Correspondingly, the observed latency range in the young and middle aged subjects, indicates the presence of both subcortical (~ 5-20 ms) and cortical contributions (~ 20-40 ms) to the response. Importantly, the more limited significant latency range for older subjects, i.e. from 14 and 19 ms, suggests a loss of cortical contributions to the response at older age.

By plotting the TRFs on a topoplot, the spatial properties of the responses can be studied. The spatial response patterns of the young subjects indicated a combination of central activity and right lateralized posterior-temporal activity, matching earlier reported findings for the same stimulus in a different dataset (Van Canneyt et al., 2020b). In our previous work, we hypothesized that the central activity may be generated mostly by sources in the brainstem, including the inferior colliculus, the cochlear nucleus and the thalamus. The right lateralized posterior temporal activity may stem from the right primary auditory cortex. The observed spatial patterns therefore suggest the presence of both subcortical and cortical response components for young subjects. Activity in both regions reduced with advancing age, with the posterior-temporal activity vanishing completely for the oldest age group. Again, this indicates a loss of cortical contributions to the response at older age. However, it is important to note that our methods only provide a rough estimate of the spatial distribution of the response and that for true source analysis, different methods are better suited (e.g. Farahani et al., 2020).

Through animal studies and clever experimental design, researchers have identified possible anatomical and physiological mechanisms that underlie the effects of age on the auditory system. Evidence suggests that reduced levels of inhibitory neurotransmitters (Caspary et al., 1995; Hughes et al., 2010), temporal jitter (Pichora-Fuller et al., 2007), prolonged neural recovery and increased neural noise may interfere with neural synchronization in the auditory system of older adults. These age-related effects disturb the precise neural coding of temporal auditory information and likely occur for both subcortical and cortical neurons. For subcortical neurons, the unique inhibitory circuitry that allows for extremely fast and precise temporal coding may falter with advancing age. This is evidenced by studies with high-frequency fine structure FFRs (for which cortical contributions are absent), which have found that higher frequencies (~ 1000 Hz) (Clinard et al., 2010) and fast sweeping frequencies (~ 1333-6667 Hz/s) (Clinard and Cotter, 2015) evoke smaller subcortical responses in older vs. younger adults. Remarkably, Clinard et al. (2010) showed that lower frequency responses (~ 500 Hz) were relatively unaffected by age, indicating that age may shift the maximum frequency that is reliably represented in the subcortical neurons, rather than equally reducing the response at any frequency. For cortical neurons, a similar shift in the maximum phase-lockable (modulation) frequency may happen. Since this frequency threshold for cortical neurons is already relatively low in young subjects (~ 150 Hz), it is possible that it may shift below the f0 range for subjects of advanced age, preventing cortical sources from contributing to the f0 response. Our results match this hypothesis: the limited range of significant latencies and the missing posterior temporal activity for the older subjects indicate a decrease in, and even absence of, cortical contributions to the f0 response with older age.

When discussing the effect of age on neural phase-locking responses, it is important to take the modulation or f0 frequency of the evoking stimulus into account. At modulation or f0 frequencies above 150 Hz, where only subcortical sources are at play, the phase-locking response is likely to decrease with age, especially for dynamic stimuli of higher frequency. At modulation or f0 frequencies between 50 and 150 Hz, were both subcortical and cortical sources are at play, both components decrease with age and the cortical contribution may be completely eliminated at older ages. Below 50 Hz, cortical sources dominate the response and curiously, evidence points towards an *increase* in response strength with advancing age. For example, Goossens et al. (2016) and Farahani et al. (2020) describe a decrease in ASSR response strength for higher frequencies (~ 80 Hz), but an *increase* in ASSR response strength for lower frequencies (< 50 Hz) with advancing age. Moreover, envelope tracking responses (typically < 30 Hz) have also been found to *increase* with advancing age (Presacco et al., 2016; Decruy et al., 2019)). These results indicate that ‘lower’ frequency auditory information is still properly phase-locked to by cortical sources, and is in fact *better* represented in the cortical activity of older adults.

The age-induced response enhancement for lower frequency auditory information has been attributed to a central gain mechanism (Parthasarathy et al., 2019) that is set into motion by reduced afferent input. The cochlear synaptopathy that commonly occurs with advancing age (Parthasarathy and Kujawa, 2018), causes auditory neurons further along the auditory pathway to receive reduced input. Through corticofugal adaptive processes, the auditory system may compensate for this by reducing inhibitory neurotransmitters (De Villers-Sidani et al., 2010). This adaptation process increases excitation in the neurons, as early as the cochlear nucleus (Wang et al., 2009), and enhances the neural response. However, the reduced inhibition is detrimental for temporal precision and response selectivity in the auditory pathway, leading to imprecise temporal coding of higher-frequency speech features (e.g. the f0) (Chambers et al., 2016). Thus, the mechanism may provide larger responses for low-frequency speech features (e.g. the envelope < 50 Hz), but it also leads to poorer response for high-frequency speech features (e.g. the f0). This explains why Decruy et al. (2019) found that advancing age increased the envelope-tracking response, whereas in the present study, with the same EEG data, we found that age decreased the f0-tracking response.

### 4.2. The effect of hearing loss on the f0 response

In a second step, we investigated the effect of hearing loss on the f0 response. Subjects with a hearing impairment had significantly larger response strength compared to age-matched normal-hearing controls, indicating a f0 response enhancement with hearing loss. These findings contradict the result of some earlier FFR studies that show that hearing loss either does not affect (Presacco et al., 2019; Roque et al., 2019), or decreases the response (Ananthakrishnan et al., 2016; Hao et al., 2018). However, as pointed out by Anderson and Karawani (2020), the results of these studies may be biased by an age effect, since the considered hearing-impaired subjects were all of older age. Since age reduces the f0 response, any enhancing effects of hearing loss may have been reduced or cancelled out by the decreasing effect of older age. In contrast, Anderson et al. (2013a) and Goossens et al. (2019) considered young, middle-aged and older hearing impaired subjects, as well as age-matched normal hearing controls, and found larger responses to the f0 (or modulation frequency in the f0 range) for subjects with a hearing impairment than without, matching the present results. In fact, Goossens et al. (2019) found no effect of hearing impairment in the oldest adults, supporting the theory of an interaction between age-related reduction and hearing-loss related enhancement of the response.

The TRF analysis in forward modelling allowed us to study the temporal properties of the response. The average TRF of the hearing-impaired subjects was significantly different from the normal hearing controls for latencies between 37.8 and 50 ms. More specifically, the subjects with a hearing impairment displayed large and dominant activity at around 45 ms latency, which was absent in age-matched normal hearing controls. Moreover, the amplitude of this response activity was significantly related to the PTA of the subjects, with larger response activity corresponding to more severe hearing loss. The latency suggests that this additional activity is cortical, and it occurs later than the cortical response contributions observed in young normal hearing subjects. From the topoplots, we know that this activity is generated centrally as well as widely-spread throughout occipital and parietal regions.

As discussed earlier, similar response enhancement has been observed for envelope responses in subjects of advancing age. Prior studies have also found increased envelope-tracking responses for subjects with a hearing impairment (Decruy et al., 2020; Fuglsang et al., 2020). In both cases, it has been theorized that the reduced afferent input (due to age or hearing loss) activates a central gain mechanism, which increases neural excitability and boosts response amplitudes (Anderson and Karawani, 2020). However, it is unlikely that this mechanism also explains the hearing loss related enhancement observed for the f0 response in the present study. As explained earlier, the central gain mechanism is detrimental for phase-locked responses to frequencies in the f0-range and actively decreases the f0 response. Thus, even though the central gain mechanism takes place in subjects with a hearing impairment, decreasing response amplitudes, there has to be a second mechanism that boosts the f0 response.

Even though both age and hearing loss are related to anato-physiological disturbances in the auditory periphery, the extent of the damage is likely greater in subjects with a diagnosable hearing loss. With this in mind, we may speculate about the underlying mechanism for the observed response enhancement. A first important aspect to consider is listening effort. Despite the fact that the hearing-impaired subjects listened to the story in an aided way and reported good comprehension, they likely put more effort in to fully understand it than normal-hearing subjects. In contrast with long-standing belief, recent findings suggests that ‘subcortical’ responses are affected by attention (Etard et al., 2019; Holmes et al., 2018), so greater listening effort may have led to exaggerated neural responses. Moreover, increased listening effort has often been associated with increased activity in the prefrontal cortex, premotor cortex, and the cingulo-opercular network (Peelle, 2018). These neural sources are involved in listener’s attention, articulatory motor planning and verbal short-term memory (Peelle et al., 2010), and may correspond to the observed central response location. It is an interesting challenge for future research to quantify the relation between listening effort and the f0 response, but as discussed in Decruy et al. (2020), various measures for listening effort exist and their relative reliability is under debate.

A second factor, that is likely more important than the augmented listening effort during the experiment itself, is the long-term speech perception difficulties experienced by hearing impaired subjects in daily life. The subjects likely have dealt with long periods of inadequate auditory perception. Even though hearing aids can increase audibility, they cannot restore the decreased temporal and spectral resolution of auditory processing. As a result, hearing impaired subjects struggle with speech understanding in noise on a daily basis. It is therefore not surprising that a significant amount of cortical reorganisation takes place in their brain: several studies have found evidence for the recruitment of additional neural resources to aid with speech comprehension when the acoustic signal is degraded due to hearing loss (Peelle et al., 2010; Du et al., 2016; Campbell and Sharma, 2013; Cardin, 2016). The wide-spread activity in the topoplots of hearing impaired subjects in figure 4 supports the theory that additional neural resources contribute to the f0 response in subjects with a hearing impairment. More specifically, it seems that the same structures that become active with increased listening effort, i.e. the prefrontal cortex, the premotor cortex and the cingulo-opercular network, may become a fully integrated part of the auditory processing network in subjects with hearing impairment (Peelle et al., 2010; Du et al., 2016; Peelle, 2018). Both the cingulo-opercular network and the premotor cortex could match with the central activity observed in the topoplots, however more precise source analysis is required to confirm this theory. Besides central activity, the topoplots also indicate diffuse parietal and occipital activity in subjects with a hearing impairment. Farahani et al. (2019) has identified several occipital and parietal neural sources for auditory temporal processing outside the primary auditory pathway. These contribute relatively weakly to auditory phase-locked responses in normal hearing subjects, but may become more active in subjects with a hearing impairment. The increased activity in the non-primary sources may compensate for reduced activity from the primary auditory pathway, as studies have found evidence for reduced activation and even gray matter atrophy in the primary auditory cortex of hearing impaired subjects (Peelle et al., 2011; Campbell and Sharma, 2013; Cardin, 2016).

Besides these two factors, other unknown factors may be at play here and further research is needed to pinpoint the exact mechanism underlying the enhanced f0 responses. One important consideration is that in order to contribute to the f0 response, a neural source needs to be able to phase-lock to f0 frequencies. It is known that some cortical sources can respond up to 150 Hz, but as discussed in the previous section, this frequency limit seems to decrease with age due to the central gain mechanism. With this in mind, the present results suggest two things: 1) the additional cortical sources that are recruited in subjects with a hearing impairment have high enough temporal precision to phase-lock at f0 frequencies and 2) they have not been affected by the interfering effects of the central gain mechanism. This might be because these additional resources have not experienced a reduction in afferent input. Another important remark is that the f0 response is highly dependent on voice characteristics (Van Canneyt et al., 2020b) and the present study only considered a low-frequency male voice. It is likely that a female-narrated story with higher and more variable f0, will evoke less cortical responses and the enhancing effect of hearing loss may therefore be reduced. Further research with more stimuli is required to confirm this hypothesis.

### 4.3. Clinical applications

The f0 response is an interesting measure for clinical practice because it is objective, relatively fast and cheap. Moreover, it is quite pleasant for the participant: listening to a story is a positive experience that is familiar, even for very young children. The results of this study indicate that the f0-response can detect age-related auditory deficits, even in subjects with a clinically-normal audiogram. This may be useful to help the large amount of patients with a normal audiogram who complain about supra-threshold hearing deficits, e.g. “being able to hear that someone is speaking but not being able to understand what they say”. Moreover, the f0 response may have clinical potential for patients with diagnosable hearing loss as well. We found that a larger f0 response was significantly related to a higher degrees of hearing loss, suggesting that the f0 response may used as an objective measure for hearing loss. In addition to being related to the degree of hearing loss, which is also true for the ABR, the f0 response could provide information about the cortical compensation mechanisms a patient has developed and therefore guide the rehabilitation strategy (Anderson et al., 2013b). Further research is needed to explore the valorisation of the f0 measure in clinical practice.

## 5. Conclusion

In this study we investigated the effects of age and hearing loss on the f0 response measured with EEG. The results indicated that response strength decreased with advancing age, but increased with hearing loss. The reduction in response strength with age is likely a side-effect of a central gain mechanism. This mechanism reduces inhibitory neural processes, which increases phase-locking capacity to low-frequency features (like the envelope) but reduces phase-locking ability to higher frequency features (like the f0). The response enhancement with hearing impaired subjects is likely the result of the recruitment of additional neural sources into the auditory processing network to aid with the perception of degraded speech.

## 6. Acknowledgements

Authors would like to thank Lien Decruy and Jonas Vanthornhout for collecting the dataset used in this study. They were assisted in data collection by Elien Van den Borre, Melissa Schoubben, Annelies Devesse and Sam Denys. This research was funded by TBM-project LUISTER (T002216N) from the Research Foundation Flanders (FWO) and also jointly by Cochlear Ltd. and Flanders Innovation & Entrepreneurship (formerly IWT), project 50432. Additionally, this project has received funding from the European Research Council under the European Unions Horizon 2020 research and innovation programme (grant agreement No. 637424, ERC starting grant to Tom Francart). The first author, Jana Van Canneyt, is supported by a PhD grant for Strategic Basic research by the Research Foundation Flanders (FWO), project number 1S83618N. Finally, the research is carried out with support from a Wellcome Trust Collaborative Award in Science RG91976 to Dr. Bob Carlyon and Jan Wouters, and with support from Flanders Innovation & Entrepreneurship through the VLAIO research grant HBC.2019.2373 with Cochlear. There are no conflicts of interest, financial, or otherwise.

## Abbreviations

ABR: auditory brainstem response
ASSR: auditory steady-state response
CAEP: cortical auditory evoked potential
EEG: electroencephalogram
FFR: frequency following response
f0: fundamental frequency of the voice
PTA: pure tone average
TRF: temporal response function

